# PyEcoLib: a python library for simulating *E. coli* stochastic size dynamics

**DOI:** 10.1101/2020.09.29.319152

**Authors:** Camilo Blanco, Cesar Nieto, Cesar Vargas, Juan Pedraza

**Affiliations:** Deparment of Mathematics and engineering. Fundacion Universitaria Konrad Lorenz. Bogota. Colombia; Department of Physics. Universidad de los Andes. Bogota. Colombia; Corporacion Colombiana de Investigacion Agropecuaria. Mosquera. Colombia

## Abstract

Recent studies describe bacterial division as a jump process triggered when it reaches a fixed number of stochastic discrete events at a rate depending on the cell-size. This theoretical approach enabled the computation of stochastic cell-size transient dynamics with arbitrary precision, with the possibility of being coupled to other continuous processes as gene expression. Here we synthesize most of this theory in the tool PyEcoLib, a python-based library to estimate bacterial cell size stochastic dynamics including continuous growth and division events. In this library, we include examples predicting statistical properties seen in experiments.

---

High-throughput measurements of microorganisms such as bacteria [1–3], yeast [4,5], and archaea [6] have shown insights of the stochastic nature of cell division dynamics and its consequences for cell physiology, shape, and gene product concentration [7–9].

Depending on how stochastic models answer the question about the mechanism used by bacteria to decide when to divide, these models can be divided in two main groups: Discrete stochastic maps (DSMs) and Continuous Rate Models (CRMs) [10].DSMs, traditionally the most used, are based on the idea that at a phenomenological, coarse-grained level, the division strategy is a map that takes cell size at birth *s_b_* to a targeted cell size at division *s_d_* trough a deterministic function *s_d_* = *f*(*s_b_*) plus stochastic fluctuations that have to be fitted from experiments [11].

In contrast to the simple description given by DSMs approach, continuous rate models (CRMs) explain not only these mapping but the entire cell cycle dynamics. Recent experiments and mathematical modeling have suggested size-dependent accumulation of FtsZ up to a critical threshold to be a putative biophysical mechanism triggering division [12,13]. Mathematically, this FtsZ accumulation can be modelled as the occurrence of some division steps, each happening at an associated continuous rate *h* that sets the probability of step occurrence into an infinitesimal time interval [14,15].

In this article, we synthesise the known CRMs describing cell division dynamics developing PyEcoLib, a python based library that can be used to estimate the cell size dynamics considering stochasticiy in division steps resembling most of the known properties of bacterial division: the division strategies [10,16], the known distributions in division times [1], fluctuations in both growth rate and in septal position [17,18]. There are two main tools in this library: first, a numerical estimator that, using the Finite State Projection algorithm [19], solves numerically the master equation describing the model [20, 21] and, second, a simulator that uses the standard Stochastic Simulation Algorithm [22] to generate division times distributed according our theory predictions. This simulator can be easily incorporated to any python based simulation scripts to model gene expression including size effects in bacterial physiology.

The main idea behind the division process, as we explained before [15], consists on the splitting being triggered once some specific number of division steps happening in a stochastic way at a rate considered to be a function of the cell size [15, 23]. Depending on the distribution of the hitting times of this division steps process, the distribution of size at division can be estimated considering exponential growth as well.

Some known properties of cell division have already observed experimentally [1,15,16]. One of these properties is the division strategy which is traditionally explained in terms of DSMs. In optimal growth conditions, the cell grows adding, on average, a fixed size since the last division event [1]. Thus, the added size Δ does not depend on *s_d_*. This strategy, colloquially known as the *adder,* can be obtained, in terms of the CRMs, if the rate of occurrence of the division steps is considered to be proportional to the cell size [15, 24].

Other division strategies can be found in some rod-shaped organisms like slow growing *E. coli,* yeast *E. pombi* [4] and *c. crescentus* [2] include the timer-like, where the added size is positively correlated with the size at birth or the sizer-like where this correlation is negative, can be obtained, in the same way, if the division rate is assumed to be a power law of the cell-size.

Additional properties of cell division can be studied using PyEcoLib. For instance some stochastic fluctuations in the position of the septum ring–The place where bacteria get split–, were found [17]. Although, on average, cell get split on a half, in some growth conditions, the stochastic fluctuations around this value can be as high as 5% [2, 17].

Cell-to-cell variability in growth rate [25, 26], can also be modelled using PyEcoLib. This fluctuation can be as high as 10% [1]. This growth rate is defined at the beginning of each cell cycle and remains constant over the entire cycle with no additional stochastic fluctuations and with no correlation with past cycles.

The library can be found in our repository ^1^ and can also be installed using the Python Package Index (pip install PyEcoLib).

How the fluctuations affect the size dynamics is studied using PyEcolib and plotted in Figure 1. Three different scenarios were considered. In Figure 1 a. we plotted the mean size dynamics obtained from a simulation of 5000 cells starting from their most recent division. We assumed that they have the same growth rate (noiseless) and split in a perfectly symmetric way. After division, it is assumed that the population number remains constant: one cell of the offspring is discarded.

**Figure 1:**
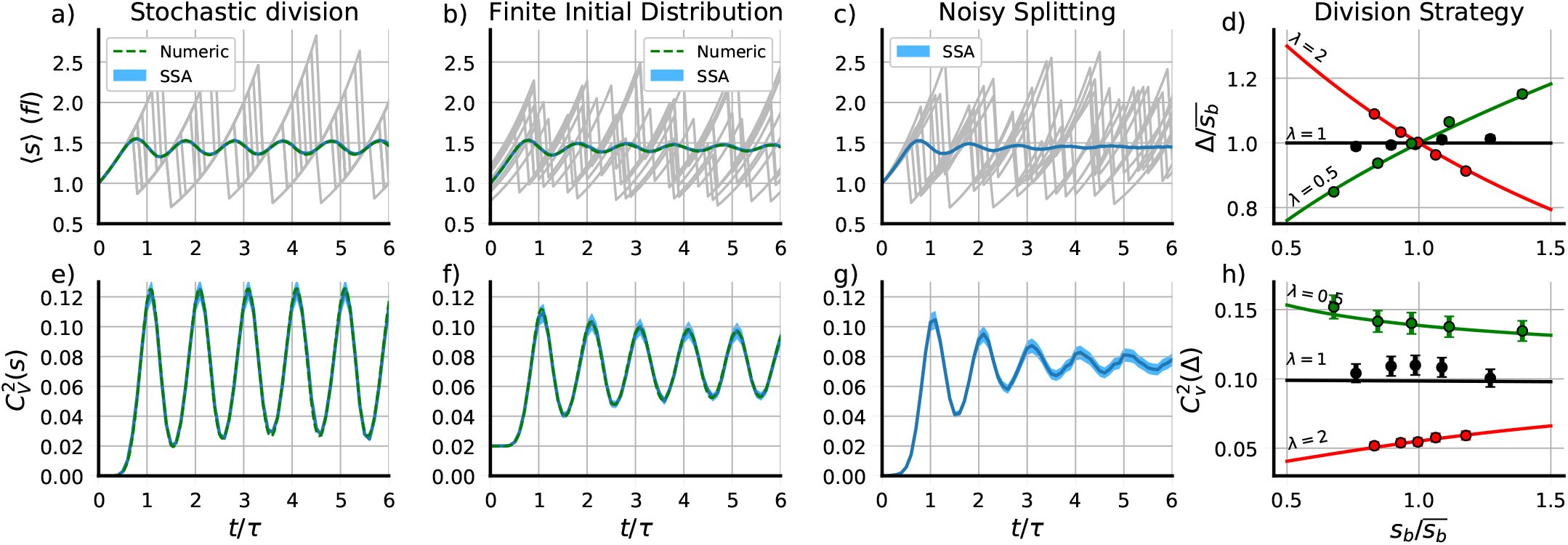
Main properties of bacterial cell division explored using PyEcoLib. a) Mean cell size 〈*s*〉 and e) Its variability 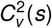 along the time considering only stochastic division. b) Mean cell size and f) Its variability along the time considering both stochastic division and an initial size distribution with finite variance. c) Mean cell size and g) Its variability along the time considering stochastic division and noise in cell-to-cell growth rate and septal position. d) mean added size Δ vs the size at birth *s_b_* and h) the fluctuations 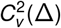 vs *s_b_* for different division strategies. Timer-like (λ = 0.5), adder (λ = 1) and sizer-like (λ = 2). Simulations (dots) and numerical estimations (lines) are shown. *M* =10 division steps were considered in all cases.

**Figure 2:**
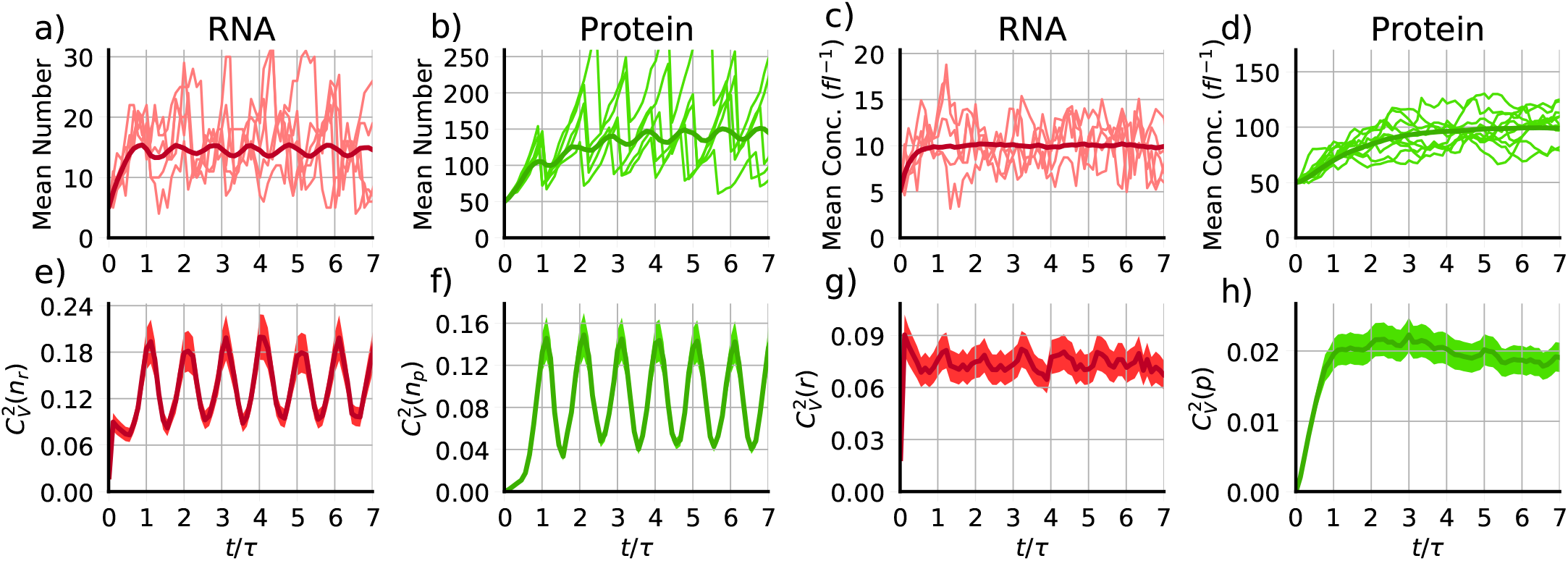
Dynamics of the main properties of gene expression for a constitutive gene. a) Mean RNA number and e) its fluctuations along the time. b) Mean protein number and f) its fluctuations along the time. c) Mean RNA concentration and g) its fluctuations along the time. d) Mean protein concentration and h) its fluctuations along the time. Parameters were set such as the mean added size was 〈Δ〉 = 1*fl*, the mean RNA concentration was 10 (*fl*^−1^) and the mean protein concentration was 100 (*fl*^−1^).

PyEcoLib can present this size dynamics using simulations and numerical methods. For simulation, PyEcoLib generates random times distributed from the cumulative function found in past studies [**?**]. Numerical methods are based on the integration of the associated master equation defining the process [21]. The dynamics of the mean size 〈*s*〉 is presented in Figure 1.a. together with ten examples of cell cycles plotted on the background to observe how variable the single-cell histories are. The dynamics of this variability of size distribution, quantified by the coefficient of variation 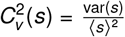, is also shown in Figure 1.e. As main effect observed, we can highlight the oscillations in both 〈*s*〉 and 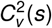 with period equal to *τ*, the doubling time. The oscillations in the 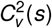 present their peaks just when bacteria are dividing on average and their valleys when bacteria are growing.

In figure 1. b., 〈*s*〉, corresponds to cells with an initial distribution with finite variance (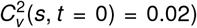. While simulations were modified by simply using random initial sizes, numerical estimation was done by performing a convolution of the solutions over the initial size distribution. Dynamics on cell size variability 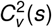 are also presented in Figure 1. f. Similar oscillations were found in both the mean and the variability but with less amplitude than the first case.

A third scenario consists on the assumption that bacteria do not split perfectly on a half but in a random variable centered on 0.5 and with a given variability 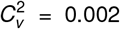. Noise in cell-to-cell growth-rate was also considered and set to 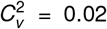. The dynamics on 〈*s*〉 was presented in Figure 1.c and the cellsize variability, presented as well in Figure 1.g. The numerical approach were not doable and thus not presented.

PyEcoLib is also able to estimate the division strategy, in terms of DSMs, using both simulations and numerical estimations. In Figure 1 d. we present the mean added size *δ* as function of the mean size at birth *s_b_* for different strategies (Different slopes in added size vs size at birth) that can be obtained changing a parameter λ. These lambda were chosen to represent three of the most important division strategies: timer-like (0 < λ < 1, where we choose λ = 0.5) with its characteristic positive slope on Δ vs *s_b_*, Adder (λ = 1) with no correlation between Δ and *s_b_* and sizer-like (1 < λ < ∞, where we choose λ = 2) with a negative slope in Δ vs *s_b_*. Fluctuations over these trends are also shown in figure 1 h. where it can be seen that sizer-like shows positive correlation in 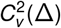 vs *s_b_,* adder strategy shows no-correlation and timer-like shows a negative correlation.

The dependence of the cell-size dynamics on gene expression can also studied. This process, gene expression, is traditionally taken as a first order reaction dependent only on the concentration of typical reactants like gene-loci, RNA and proteins [27–29]. When size dynamics is considered, as we explained in recent studies [20], differential equations in molecule concentration can be transformed to molecule number inside the cells which are growing [20, 30].

During division, whose times are estimated using PyEcoLib, we propose that these molecules are segregated to each descendant cell following a binomial distribution with parameter 0.5 but this parameter can be changed using PyEcolib as well if not-symmetric partition is considered.

We set the parameters such as the steady size at birth was 1 fl [1], the time was measured in units of doubling time *τ* = ln(2)/*μ* with *μ* being the the elongation rate such as the cell grows following 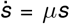. Using PyEcoLib, we set this doubling time to be τ = 18 min ant the normalization was made later. The mean RNA concentration was 10 molecules/fl and active degradation *γ_r_* = 5*μ* per molecule was considered. Protein concentration was set to have a steady value of 100 proteins/fl and no active degradation. Initial conditions was set such as they started at fixed in one half of their steady concentration. Given this, the starting

In Figure 3 a) we observe the dynamics of the mean RNA number *n_r_* inside the cells with some single-cell trajectories presented on the background. Fluctuations around this trend are quantified in Figure 3 e. Protein number dynamics (*n_p_*) (Fig 3 b.) and its fluctuations (Fig 3. f.) are also presented. We observe oscillations in these trends in similar way to the size dynamics showing that these molecule numbers are very correlated to the size.

Dynamics of RNA mean concentration 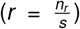 is presented in Figure 3.a) and stochastic fluctuations over this trend are presented in Figure 3.g. Finally, dynamics on mean protein concentration (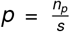, Fig 3.d) and its fluctuations (Fig 3.h.) are presented as well. We can highlight how the concentration of these molecules show almost no oscillations.

In this article we presented PyEcoLib, a python based library using the current continuous rate theory of bacterial cell division models [15,21] which consider the division as a time-continuous stochastic jump process triggered upon the occurrence of a fixed division steps. We use PyEcoLib to estimate the stochastic dynamics of the cell size for a population of constant number. This constant number can be obtained if one of the bacterial descendants is discarded and the other one is still tracked as happens in micro-fluid Mother machine experiments [1]. Some attempts to estimate cell size dynamics in populations of growing number have been already done using both numerical methods and stochastic simulation [23,31]. However, since the number of cells in these populations grows exponentially on time, these simulations can become unstable when this population number is high. In future versions of the library we think to incorporate this tool.

Additionally to estimate the size dynamics, some possibilities of this library include: Simulating most of the division strategies found in E. coli: timer-like, adder and sizer-like [10,16]. Estimate the distribution of division times, size at division and size at birth in a population of constant number. The library can be coupled to stochastic simulation of gene expression with reactions happening at arbitrary distributed times. Noise in both septal ring placing and cell-to-cell growth rate can be considered [17]. An arbitrary size distribution at the starting time can be also considered, however an arbitrary starting distribution of division steps is not already considered and all the cells must start with zero division steps.

As is shown in Figure 1, oscillations in both, the mean size 〈*s*〉 and 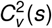, were found. When only stochasticity on division steps is considered, these oscillations are maintained over a arbitrary long period of time. These oscillations are also found but with less amplitude when an initial size distribution with finite variance is considered and are now damped when other sources of noise like the cell-to-cell growth rate variability and septal position are added. This phenomenon can be explained considering the maintenance of correlation between cell size along multiple cell cycles: Cells get born in a given time *t* are very likely to divide again after a time *t* + *nτ* with *n* and arbitrary integer, this correlation decays to zero, however, where the other noises are added. The larger the time in the future, the more uncertain will be time that cell will got split.

Including cell-size stochasticity in gene expression can be an important tool to understand the origin of the fluctuations in molecule concentration. Some efforts have already been done to understand these effects in simple regulatory networks [20, 32–35] but the use of PyEcoLib in more complex gene regulatory architectures, can improve the understanding the effects not only of the stochastic in division but the effects of other complex process in division such as the division strategy, the noise in growth rate and the asymmetric cell splitting.

## Footnotes

1 https://github.com/SystemsBiologyUniandes/PyEcoLib

## References

[1] S. Taheri-Araghi, S. Bradde, J. T. Sauls, N. S. Hill, P. A. Levin, J. Paulsson, M. Vergassola, and S. Jun, “Cell-size control and homeostasis in bacteria,” Current biology, vol. 25, no. 3, pp. 385–391, 2015.

[2] S. Iyer-Biswas, C. S. Wright, J. T. Henry, K. Lo, S. Burov, Y. Lin, G. E. Crooks, S. Crosson, A. R. Dinner, and N. F. Scherer, “Scaling laws governing stochastic growth and division of single bacterial cells,” Proceedings of the National Academy of Sciences, vol. 111, no. 45, pp. 15912–15917, 2014.

[3] M. Campos, I. V. Surovtsev, S. Kato, A. Paintdakhi, B. Beltran, S. E. Ebmeier, and C. Jacobs-Wagner, “A constant size extension drives bacterial cell size homeostasis,” Cell, vol. 159, no. 6, pp. 1433–1446, 2014.

[4] J.-B. Nobs and S. J. Maerkl, “Long-term single cell analysis of s. pombe on a microfluidic microchemostat array,” PloS one, vol. 9, no. 4, p. e93466, 2014.

[5] D. Chandler-Brown, K. M. Schmoller, Y. Winetraub, and J. M. Skotheim, “The adder phenomenon emerges from independent control of pre-and post-start phases of the budding yeast cell cycle,” Current Biology, vol. 27, no. 18, pp. 2774–2783, 2017.

[6] Y.-J. Eun, P.-Y. Ho, M. Kim, S. LaRussa, L. Robert, L. D. Renner, A. Schmid, E. Garner, and A. Amir, “Archaeal cells share common size control with bacteria despite noisier growth and division,” Nature microbiology, vol. 3, no. 2, p. 148, 2018.

[7] L. Willis and K. C. Huang, “Sizing up the bacterial cell cycle,” Nature Reviews Microbiology, vol. 15, no. 10, pp. 606–620, 2017.

[8] A. Raj and A. Van Oudenaarden, “Nature, nurture, or chance: stochastic gene expression and its consequences,” Cell, vol. 135, no. 2, pp. 216–226, 2008.

[9] W. J. Blake, G. Balázsi, M. A. Kohanski, F. J. Isaacs, K. F. Murphy, Y. Kuang, C. R. Cantor, D. R. Walt, and J. J. Collins, “Phenotypic consequences of promoter-mediated transcriptional noise,” Molecular cell, vol. 24, no. 6, pp. 853–865, 2006.

[10] P.-Y. Ho, J. Lin, and A. Amir, “Modeling cell size regulation: From single-cell-level statistics to molecular mechanisms and population-level effects,” Annual review of biophysics, vol. 47, pp. 251–271, 2018.

[11] Y. Tanouchi, A. Pai, H. Park, S. Huang, R. Stamatov, N. E. Buchler, and L. You, “A noisy linear map underlies oscillations in cell size and gene expression in bacteria,” Nature, vol. 523, no. 7560, pp. 357–360, 2015.

[12] F. Si, G. Le Treut, J. T. Sauls, S. Vadia, P. A. Levin, and S. Jun, “Mechanistic origin of cell-size control and homeostasis in bacteria,” Current Biology, vol. 29, no. 11, pp. 1760–1770, 2019.

[13] K. Sekar, R. Rusconi, J. T. Sauls, T. Fuhrer, E. Noor, J. Nguyen, V. I. Fernandez, M. F. Buffing, M. Berney, S. Jun, et al., “Synthesis and degradation of ftsz quantitatively predict the first cell division in starved bacteria,” Molecular systems biology, vol. 14, no. 11, p. e8623, 2018.

[14] K. R. Ghusinga, C. A. Vargas-Garcia, and A. Singh, “A mechanistic stochastic framework for regulating bacterial cell division,” Scientific reports, vol. 6, p. 30229, 2016.

[15] C. Nieto, J. Arias-Castro, C. Sánchez, C. Vargas-García, and J. M. Pedraza, “Unification of cell division control strategies through continuous rate models,” Physical Review E, vol. 101, no. 2, p. 022401, 2020.

[16] J. T. Sauls, D. Li, and S. Jun, “Adder and a coarse-grained approach to cell size homeostasis in bacteria,” Current opinion in cell biology, vol. 38, pp. 38–44, 2016.

[17] S. Modi, C. A. Vargas-Garcia, K. R. Ghusinga, and A. Singh, “Analysis of noise mechanisms in cell-size control,” Biophysical journal, vol. 112, no. 11, pp. 2408–2418, 2017.

[18] A.-C. Chien, N. S. Hill, and P. A. Levin, “Cell size control in bacteria,” Current biology, vol. 22, no. 9, pp. R340–R349, 2012.

[19] B. Munsky and M. Khammash, “The finite state projection algorithm for the solution of the chemical master equation,” The Journal of chemical physics, vol. 124, no. 4, p. 044104, 2006.

[20] C. Nieto-Acuña, J. C. Arias-Castro, C. Vargas-García, C. Sánchez, and J. M. Pedraza, “Correlation between protein concentration and bacterial cell size can reveal mechanisms of gene expression,” Physical Biology, vol. 17, no. 4, p. 045002, 2020.

[21] C. A. Nieto-Acuna, C. A. Vargas-Garcia, A. Singh, and J. M. Pedraza, “Efficient computation of stochastic cell-size transient dynamics,” BMC bioinformatics, vol. 20, no. 23, pp. 1–6, 2019.

[22] D. T. Gillespie, “A general method for numerically simulating the stochastic time evolution of coupled chemical reactions,” Journal of computational physics, vol. 22, no. 4, pp. 403–434, 1976.

[23] N. Totis, C. Nieto, A. Küper, C. Vargas-García, A. Singh, and S. Waldherr, “A population-based approach to study the effects of growth and division rates on the dynamics of cell size statistics,” IEEE Control Systems Letters, vol. 5, no. 2, pp. 725–730, 2020.

[24] C. A. Vargas-García and A. Singh, “Elucidating cell size control mechanisms with stochastic hybrid systems,” in 2018 IEEE Conference on Decision and Control (CDC), pp. 4366–4371, IEEE, 2018.

[25] D. J. Kiviet, P. Nghe, N. Walker, S. Boulineau, V. Sunderlikova, and S. J. Tans, “Stochasticity of metabolism and growth at the single-cell level,” Nature, vol. 514, no. 7522, pp. 376–379, 2014.

[26] S. Vadia and P. A. Levin, “Growth rate and cell size: a re-examination of the growth law,” Current opinion in microbiology, vol. 24, pp. 96–103, 2015.

[27] J. Paulsson, “Models of stochastic gene expression,” Physics of life reviews, vol. 2, no. 2, pp. 157–175, 2005.

[28] P. Robert et al., “Mathematical models of gene expression,” Probability Surveys, vol. 16, pp. 277–332, 2019.

[29] M. Kaern, T. C. Elston, W. J. Blake, and J. J. Collins, “Stochasticity in gene expression: from theories to phenotypes,” Nature Reviews Genetics, vol. 6, no. 6, pp. 451–464, 2005.

[30] J. Jedrak, M. Kwiatkowski, and A. Ochab-Marcinek, “Exactly solvable model of gene expression in a proliferating bacterial cell population with stochastic protein bursts and protein partitioning,” Physical Review E, vol. 99, no. 4, p. 042416, 2019.

[31] P. Thomas, “Analysis of cell size homeostasis at the single-cell and population level,” Frontiers in Physics, vol. 6, p. 64, 2018.

[32] R. Perez-Carrasco, C. H. Beentjes, and R. Grima, “Effects of cell cycle variability on lineage and population measurements of mrna abundance,” BioRxiv, 2020.

[33] C. H. Beentjes, R. Perez-Carrasco, and R. Grima, “Exact solution of stochastic gene expression models with bursting, cell cycle and replication dynamics,” Physical Review E, vol. 101, no. 3, p. 032403, 2020.

[34] M. Soltani, C. A. Vargas-Garcia, D. Antunes, and A. Singh, “Intercellular variability in protein levels from stochastic expression and noisy cell cycle processes,” PLoS computational biology, vol. 12, no. 8, p. e1004972, 2016.

[35] P. Thomas, G. Terradot, V. Danos, and A. Y. Weiße, “Sources, propagation and consequences of stochasticity in cellular growth,” Nature communications, vol. 9, no. 1, pp. 1–11, 2018.

